# Dysregulation of chromatin via H3K27 methylation underpins differentiation arrest in Isocitrate dehydrogenase-mutant Acute Myeloid Leukaemia

**DOI:** 10.1101/2023.08.24.554641

**Authors:** Douglas RA Silveira, Prodromos Chatzikyriakou, Pilar Casares Aláez, Sarah Mackie, Roan Hulks, Olena Yavorska, Paulina Siejka-Zielińska, Alastair Smith, Marlen Metzner, Batchimeg Usukhbayar, Virginie Penard-Lacronique, Chun-Xiao Song, Anita K. Gandhi, Paresh Vyas, Thomas A. Milne, Skirmantas Kriaucionis, Maroof Hasan, Lynn Quek

## Abstract

Dysregulation of cellular differentiation is a hallmark of cancer. Isocitrate dehydrogenase (IDH) is commonly mutated multiple cancers including glioma, cholangiocarcinoma, lymphoma and Acute Myeloid Leukaemia (AML). Mutant IDH generates d-2-hydroxyglutarate that inhibits enzymes including Jumonji histone demethylases and TET2. Using primary human IDH2-mutant AML cells as a model, single cell RNA-seq and ATAC-seq, we demonstrated the continuum of cell states during restoration of neutrophilic differentiation to leukaemic progenitors by Enasidenib, a mutant IDH2 inhibitor. In cells which ultimately differentiate, there is co-expression of competing GATA2/RUNX3/SOX4-driven stem-progenitor and pro-differentiation EGR1/JUN/FOS programmes, followed by expression of cell cycle and terminal neutrophil programmes involving CEBP family and SPI1/PU.1. Genes upregulated during differentiation display loss of H3K27me3 in bivalent chromatin but not of H3K4me3, while downregulated genes are enriched for PRC2/EZH2 targets. In contrast to previous reports of a TET2-dependent mechanism for IDH-mutations, we observed only a modest link between promoter DNA CpG methylation and gene expression. For the first time in primary AML, we describe the lifting of differentiation block by de-repression of pro-differentiation genes through modulation of H3K27 demethylation in bivalent chromatin, and thus highlight a novel and important mechanism in how IDH mutations disrupt cell fates in cancer.

## INTRODUCTION

Disruption of cellular differentiation pathways is a hallmark of cancer, and mutations in isocitrate dehydrogenase (IDH) result in production of the oncometabolite d-2-hydroxyglutarate (d-2HG) which is implicated in dysregulation of cell fates in multiple cancers including glioma, cholangiocarcinoma chondrosarcoma, lymphoma and Acute Myeloid Leukaemia (AML) ^1^. IDH-mutant Myeloid-committed progenitors encounter differentiation block, resulting in accumulation of leukaemic progenitors and cytopenia in AML patients, whereas IDH mutations block hepatocyte differentiation to promote cholangiocarcinoma ^2^ . Small molecule inhibitors targeting mutant IDH have thus re-awakened the paradigm of differentiating agents to treat cancer. D-2HG inhibits α-ketoglutarate-dependent oxygenases, including Jumonji histone demethylases (J-KDMs) and TET2. Inhibitors of mutant IDH (mIDHi) including Enasidenib (ENA, an mIDH2i) and are now FDA-approved for treatment of AML ^3, 4^ and cholangiocarcinoma ^5^. As J-KDM and TET2 regulate transcription via histone and DNA demethylation respectively, in preclinical models, mIDHi are known to reverse DNA and histone hypermethylation. However, specific differentiation pathways, and how they may be restored in primary cancer tissues are unclear. While restoration of TET2 function has been proposed as a key mediator of the effects of IDH inhibitors through CpG demethylation, TET2 mutations are not a common event in IDH-mutant AML patients who develop resistance to IDH inhibitors, and evidence that links specific CpG demethylation to expression of differentiation pathways is lacking. Despite sharing the same oncometabolite, oncogenic mechanisms may also be dependent on cellular context. For example, IDH1-mutant gliomas show extensive CpG hypermethylation (∼19%) compared to non-mutant counterparts while CpG hypermethylation is modest in other IDH-mutant AML, melanoma and cholangiocarcinoma (∼2-4%) ^6^.

Our ability to track the process of cellular differentiation in primary AML leukaemic progenitors in timescales similar to those observed when patients are treated *in vivo*, and to interrogate molecular pathways in parallel, makes this an attractive model to better understand IDH-driven oncogenesis in human patients. Patients responding to mIDHi often develop leukocytosis associated with differentiation of AML blasts into neutrophils^4^, akin to stimulation of normal progenitors by granulocyte-colony stimulating factor^7^. We therefore hypothesise that mIDHi epigenetically re-programmes L-Prog to restore a normal progenitor state, prior to stimulation of normal differentiation pathways. By suppressing d-2HG, mIDHi may restore gene expression by disinhibition J-KDMs^8, 9^, but these demethylate both repressive and activating marks. AGI-6780, an ENA analogue, induces H3K4/9/27/36me3 demethylation in mIDH2 AML^10^, but the link between these histone changes and transcriptional programmes that mediate differentiation is unknown. Here, we used a multiomic approach to characterise the interplay between histone methylation epigenetic and modulation of transcriptional programmes associated with IDH-inhibitor induced neutrophilic differentiation of arrested leukaemic progenitors, and in reverse, revealed mechanisms by which IDH mutations cause differentiation arrest.

## RESULTS

### Mutant IDH2 AML progenitors are arrested at the granulo-monocytic/ macrophage progenitor stage of differentiation

AML is frequently characterised by expansion of CD34+ leukaemia progenitor (L-Prog) populations enriched for leukaemia stem cells. The majority of our cohort of 22 untreated mIDH2 AML patients (Supplementary Table S1) had expanded CD34+CD45RA+ progenitors, dominated by leukaemic (L)-lymphoid-primed multipotent progenitors (LMPP) and L-granulocyte-monocyte progenitors (GMP)^11^ (Supplementary Figure 1A-B and Supplementary Tables S1-2). L-GMP and L-LMPP transcriptomes differ by only 28 genes (Supplementary Figure 1E, Supplementary Table S3) and were combined as CD34+ leukaemic progenitors (L-Prog). To determine differentiation arrest, we compared gene expression of L-Prog and normal haematopoietic stem progenitors (HSPCs) derived from the data from Quek *et al*.^12^. L-Prog are most similar to normal (n-)GMP (Figure 1A), but are markedly expanded (average 111-fold) compared with normal CD34+CD45RA+ progenitors (Supplementary Table S2), indicating differentiation arrest. Compared with n-GMP, L-Prog have reduced expression of cell cycle (CC) genes, *MYC* and E2F targets, and mature neutrophil genes (*ELANE, PRTN3, MPO*); and increased expression of stem-progenitor transcription factors (TFs) (*GATA2* and *HOPX*), TP53-pathway genes and cyclin-dependent kinase inhibitors (*CDKN1A*/*1C*/*2A*/*2D*) (Figure 1B-C, Supplementary Table S4). Furthermore, L-Prog are of CC and enrichment of inflammatory signatures compared with n-GMP (Figure 1D).

**Figure 1.**
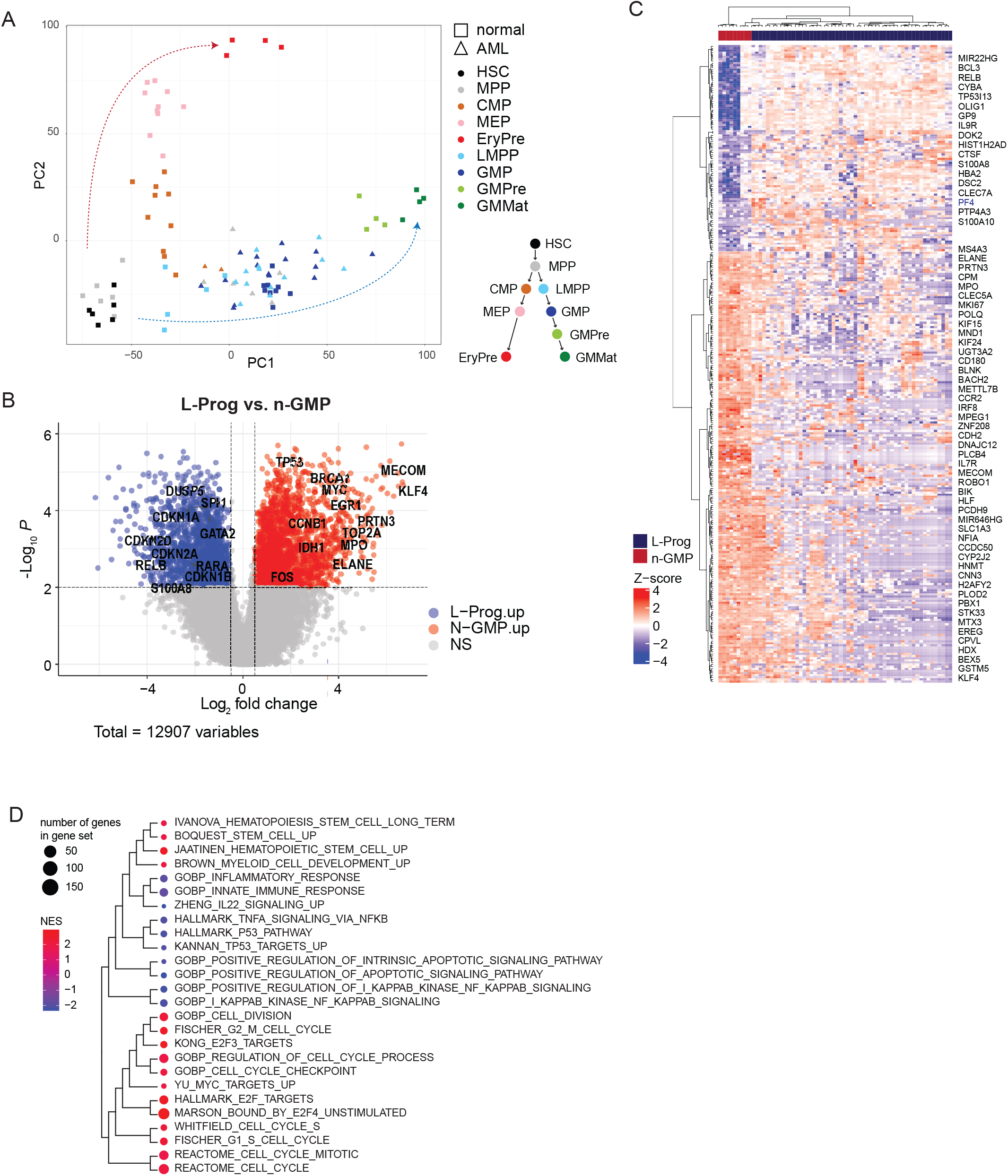
Transcriptional characterisation of IDH2-mutant AML progenitors. A) Principal component analysis (PCA) plot of flow-sorted AML progenitors (L-Prog, triangles, n=35 from 17 patients) from mIDH2 AML samples, compared with normal haematopoietic stem/progenitor/precursor and mature (HSPPM) populations (squares). Dashed arrows indicate GM (Blue) and erythroid (red) differentiation of normal haematopoietic populations. A simplified haematopoietic tree is shown for reference. See Supplementary Figure 1C for gene loadings. B) Volcano plot showing differentially expressed genes (DEG) between mIDH2 L-Prog and n-GMP. Selected genes are highlighted. Threshold of significance was set at FDR < 0.1 and a log_2_ fold change ≥0.5. C) Heatmap clustering showing z-scores of selected significant DEG (in rows) between n-GMP and mIDH2 L-Prog samples (in columns). Selected genes are annotated on the figure (full list in Supplementary Table 4). D) DEG from (C) were used for enrichment analysis. Positive normalised enrichment scores (NES) reflect genes upregulated in n-GMP, and negative NES reflect genes upregulated in L-Prog.

### Enasidenib restores normal progenitor function to AML progenitors and upregulates genes involved in normal neutrophil differentiation

We used *in vitro* culture^13^ (IVC) to study differentiation and harvested cells for assessment by flow cytometry (Figure 2A). Normal GMP from adult bone marrow proliferated and differentiated into precursors (CD34-117+, prec), CD15+ promyelocytes (prom) or CD11b+ myelomonocytic (myelomono) cells, and neutrophils (CD11b+CD15+) and monocytes (CD11b+CD14+, mono) by day (D)14 (Supplementary Figure 2A-C). In four mIDH2 L-Prog samples, ENA (1μM) treatment produced neutrophils and monocytes with varied kinetics (Figure 2B, Supplementary Figure 2D-F). Response to ENA produced more differentiated neutrophils (CD11b+/CD15+, Figure 2C) and sustained proliferation (Figure 2D) compared with DMSO. In contrast, there was no significant difference in mature s cell production, and only modest proliferation in 3 non-responsive IDH2m and 3 IDH wild-type (WT) samples (Figure 2C).

**Figure 2:**
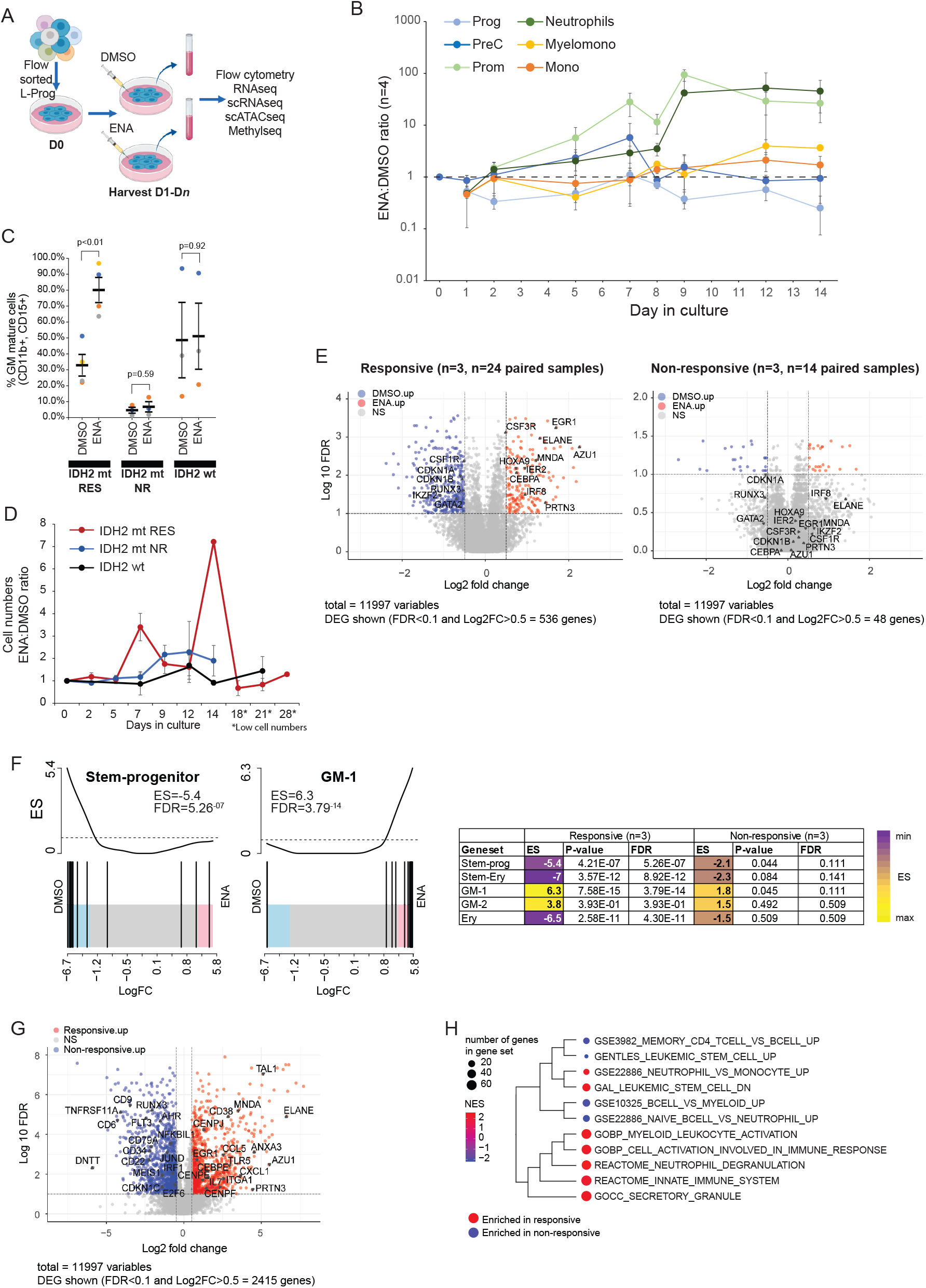
*In vitro* liquid suspension culture and differentiation of IDH2m L-Prog. A) Diagram showing *in vitro* liquid suspension culture (IVC) system and workflow. B) Output of IVC differentiation assay with L-Prog from 4 different mIDH2 AML (#3018/ 3308/ 3022/ 3162) shown as ENA: DMSO ratio, where DMSO values are normalised to 1.0 between D0 and D14 of culture. Error bars show mean +/- standard error of mean (SEM). Prog= progenitor, PreC= precursors, Prom= promyelocytes, Myelomono= myelomonocytic cells, Mono= monocytes. C) Percentage of mature GM cells (CD11b+ and/ or CD15+) produced in IVC by D14 in mIDH2 (mt. RES: Responder, n=3, NR: Non-responder, n=3) and IDH2wt L-Prog (n=3) with and without ENA (1μM). P values are from a Student ’s *t*-test, *** indicates p<0.001. Error bars denote mean +/- standard error of mean (SEM). D) Cell numbers produced in IVC expressed as ENA:DMSO ratio by L-Prog from mIDH2 and IDHwt. Other annotations and error bars are as in (C). *Marks time points where total cell numbers were low, with reduced confidence in accurary of cell counts. E) Volcano plot of DEG between DMSO and ENA-treated, from a combined analysis of flow sorted progenitor populations harvested between D1 and D12 of IVC from 3 responsive and 3 non-responsive L-Prog samples. Selected genes are highlighted. Threshold of significance was set at FDR of 0.1 and a log_2_ fold change of ≥0.5. F) Gene set enrichment (GSEA) analysis showing significant enrichment of DEG profiles of L-Prog from DMSO and ENA treated IVC (from 2E), versus gene profiles derived from normal haematopoiesis (see Supplementary Figure 2G). Two GSEA plots are presented as examples and enrichment scores (ES) and FDR values for all comparisons are shown. G) Volcano plot of DEG between ENA-treated, from a combined analysis flow-sorted L-Prog from responsive (n=3 patients) and non-responsive (n=3 patients) samples harvested between D1 and D12 of IVC. Selected genes are highlighted. Threshold of significance was set at FDR of 0.1 and a log_2_ fold change of ≥0.5. H) GSEA dendrogram showing GO gene sets with significant positive (in responsive patients) and negative (in non-responsive patients) enrichment of DEGs from (G).

To examine the transition from an arrested to un-arrested state, we performed RNAseq in DMSO or ENA-treated L-Prog at different time points. In combined pair-wise (i.e. DMSO versus ENA) analysis, ENA upregulated 174 genes, including IEGs (e.g. *EGR1, IER2*), and neutrophilic-TFs (e.g. *CEBPA*), neutrophil effector genes (*ELANE, AZU1*) and *CSF3R*. 362 genes were downregulated, including *GATA2, HOPX*, and *CDKN1A*. ENA-treated responsive L-Prog samples enrich for normal GM or neutrophil genes and are depleted of normal stem-progenitor, stem-erythroid and erythroid genes (Figure 2F, Supplementary Figure 2G, Supplementary Table S5). In contrast, there were only 48 differentially expressed genes (DEG) between ENA/DMSO-treated non-responders (Figure 2E, Supplementary Table S6). Compared with non-responsive L-Prog (Figure 2G, Supplementary Table S6), ENA-induced response specifically upregulates neutrophil genes and downregulates leukaemic stem cell (LSC) and lymphoid progenitor genes (reflecting lympho-myeloid characteristics of LSCs^14^, Figure 2H).

### Single-cell RNA-seq reveals transcriptional trajectory of Enasidenib-induced differentiation of AML progenitors

Since differentiation of L-Prog is occurs as a continuum, we performed single-cell RNA-seq (scRNA-seq) in four responsive (#3018/#3022/#3162/#3308^RES^, 26988 cells), and three non-responsive (#2917/#2755/#2814^NR^, 38523 cells) samples (Supplementary Figure 3A). Of the three non-responsive samples, #2917^NR^ IDH2-mutant (R140Q), whereas #2814^NR^ and #2755^NR^ are IDH2-WT, but are IDH1-mutant. Thus, we can compare on-target versus non-target, non-response to ENA. We harvested cells at D2, D5, D9 and D12 to capture cell types from early to late differentiation, summarised in Supplementary Figures 3A-C. Using known GM genes, we classified cells as myelomonocytic precursors/neutrophils (GM), monocytes (M) and undifferentiated (U, Supplementary Figure 3D-E). GM cells were further divided into early (EGM) or late-GM (LGM) by expression of earlier (*CST7, CLU, CYTL1, ELANE, SRGN*), or late markers (*AZU1, CSTA* and *PRTN3*). We combined differentiation states with cell cycle (CC) gene expression (G_0_, G_1_S or G2M^15^, to further annotate cells using 10 functional-CC identities (Figure 3A, Supplementary Tables S7-S8). In responsive samples, profiles were similar between DMSO and ENA at D2. At D5-D9, 22% of cells were proliferating LGM-G_2_M cells in ENA, compared with 9% in controls (OR:2.75, p<2.2e-16). At D9, non-cycling terminally differentiated (TD) neutrophils (LGM-G0) accounted for 47% of ENA-treated cells versus 13% in DMSO (OR:5.83, p<2.2e-16), but monocytes were more numerous in DMSO (8% vs. 4% at D2/D5/D9, OR:0.48, p<2.2e-16). In contrast, ENA had no significant impact on differentiation over DMSO in non-responsive samples (Figure 3B, Supplementary Figure 3F).

**Figure 3.**
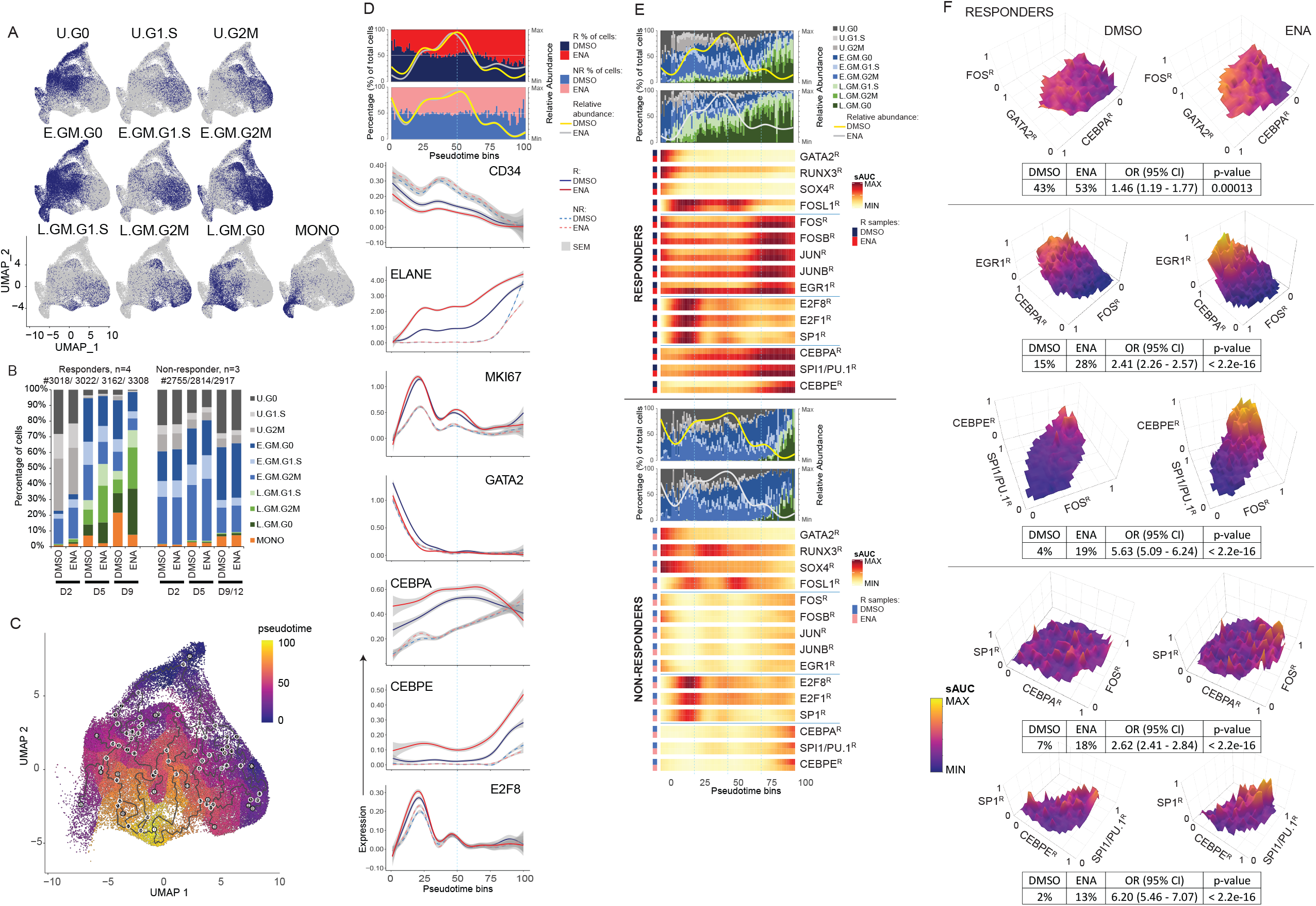
Continuum of differentiation trajectory of AML progenitors treated with Enasidenib as revealed by single cell RNA-seq. A) UMAPs showing classification of cells from four responsive and three non-responsive samples, by 10 combined functional (undifferentiated, U, early GM, EGM, late GM, LGM) and CC (G_0_, G_1_S or G_2_M) identities. Monocytes with different cell cycle states are combined as MONO. See also Supplemental Figure 3A-C. B) Bar graph showing percentage of cells with each functional and cell cycle annotation at different days in culture. Average values in combined responsive (n=4) and non-responsive (n=3) samples are shown, see also Supplemental Figure 3F for per sample data. C) Pseudotime trajectory of combined DMSO and ENA-treated cells from all seven samples. We ideintified neutrophils as the endpoint of the trajectory (cluster 13, see Supplemental Figure 3A). Pseudotime scale of 0-100 is shown. D) Upper and lower panels share the same pseudotime bin x-axis. Upper panel: Bar graph (1^st^ y-axis) showing proportion, and line graph (2^nd^ y-axis) showing relative abundance, of DMSO and ENA-treated cells in combined responsive (n=4) and non-responsive (n=3) samples versus pseudotime bins (0-100, x-axis). Colour keys for DMSO/ ENA for bars and lines are shown. Lower panels show mean expression of indicated genes that indicate differentiation and cell proliferation (CD34, ELANE, MKI67), and transcription factors (GATA2, CEBPA, CEBPE and E2F8). Data from responsive samples (n=4) and non-responsive samples (n=3) are in solid and dashed lines respectively. DMSO/ ENA conditions are coloured coded as shown. The grey area indicates standard error of mean (SEM). E) Upper and lower panels share the same pseudotime bin x-axis. Upper panel: Bar graph (1^st^ y-axis) showing average percentage of cells from DMSO and ENA treated responsive samples (n=4) with function-cell cycle annotation versus pseudotime bins. Relative abundance curves (2^nd^ y-axis) are as in (D) are also shown for ease of reference. Heatmap plots of scaled area under the curve (sAUC) values for regulons (e.g. GATA2^R^) in DMSO and ENA-treated responsive samples versus pseudotime bins. Lower panel: the same as for upper panel, but in non-responsive samples (n=3). F) 3-dimensional surface plots showing co-expression (as sAUC) of 3 regulons in DMSO and ENA-treated responsive samples (average values of 3 patients). Top panel shows in x/y/z order, GATA2^R^/CEBPA^R^/FOS^R^ with cells from pseudotime bin 1-25. Middle panels show CEBPA^R^/ FOS^R^/EGR1^R^, and FOS^R^/ SPI1/PU.1-PU.1^R^/ CEBPE^R^ with all cells. Lower panels show CEBPA^R^/FOS^R^/SP1^R^ and CEBPE^R^/SPI1/PU.1-PU.1^R^/SP1^R^ (note the different orientation of the scale 0-1 in the different plots). sAUC of the z-axis regulon is shown as a colour scale from minimum (0) to maximum (1). The percentage of cells co-expressing regulons, and the odds ratio (OR) and p-values of the comparison between DMSO and ENA are also shown alongside.

In order to focus on neutrophil differentiation for trajectory analysis, we removed monocyte clusters K7/K16, and a cluster of monocytoid blasts from one non-responsive sample (K14, Figure 3C), and designated the neutrophil cluster K13 as an endpoint. Pseudotime modelling ordered cells agnostic of response status or treatment. We confirmed pseudotime in individual samples in distribution of cells harvested from different days (Supplementary Figure 3C). Normalised cell distribution across pseudotime (DMSO vs. ENA) is in Figure 3D, alongside expression of genes that reflect differentiation (*CD34* and *ELANE*), cell proliferation (*MKI67*) and transcription factors (TFs) *CEBPA, CEBPE* (myeloid differentiation) and *E2F8* (cell cycle). The beginning and end of pseudotime represent immature L-Prog and neutrophils respectively. In responders, early pseudotime (time 0-25, enriched for U and EGM cells) is enriched for DMSO cells (61%), and the reverse is true for late pseudotime (time 76-100, 63% ENA cells). Non-responsive samples did not have significantly skewed distributions between DMSO and ENA treatment (early, 51%/49% and late, 48%/52%). Expression of *GATA2* rapidly declines during the first half of pseudotime and is significantly lower in ENA-treated responsive cells. Moreover, expression of late neutrophil differentiation genes (*ELANE/CEBPA/CEBPE*) is higher in ENA-treated responsive cells than DMSO controls, which in turn is higher than non-responsive samples. In addition, ENA induces modest increased expression of *MKI67* and *E2F8*.

To study transcriptional networks underlying these differentially expressed genes, we used SCENIC^16^ to assign highly co-expressed genes to TF motifs, to create regulatory networks (‘regulons ‘, Supplementary Table S9). Each cell may express one or more regulons (denoted by ‘^R^ ‘), and regulon and functional-CC states were correlated (Figure 3E). In early pseudotime, cells (mainly U.G0 and EGM.G0) expressed stem-progenitor regulons GATA2/RUNX3/SOX4^R^ (GRS)^R^. GRS^R^ is more rapidly downregulated with ENA in responsive samples, and GRS^R^-positive cells co-expressed higher levels of IEG/AP-1 regulons EGR1/JUN/FOS^R^ and neutrophilic-TF regulons, CEBPA/SPI1/PU.1/CEBPE^R^ during early pseudotime (Figure 3E). Increased triple co-engagement of GATA2/FOS/CEBPA^R^ in ENA versus DMSO-treated responsive samples is shown in Figure 4F (top panel). Generation of LGM cells (culminating in neutrophils) with ENA treatment correlates with sustained co-engagement of EGR1/JUN/FOS/CEBPA/SPI1/CEBPE^R^ (Figure 3E-F). Differentiation is markedly delayed with DMSO, and most cells remain EGM (precursors). Non-responsive samples had prolonged expression of GRS^R^ and minimal engagement of IEG and GM-TF regulons (Figure 3E). Expression of cell cycle regulons E2F/SP1^R^ mirrored expression of *MKI67* (Figure 3D). While the proportion of cycling cells were similar between DMSO and ENA-treated responsive samples, ENA treatment co-engaged E2F/SP1^R^ with IEG and neutrophilic-TF programmes (Figure 3E, Figure 3F, lower panel), resulting in more cycling LGM cells.

**Figure 4.**
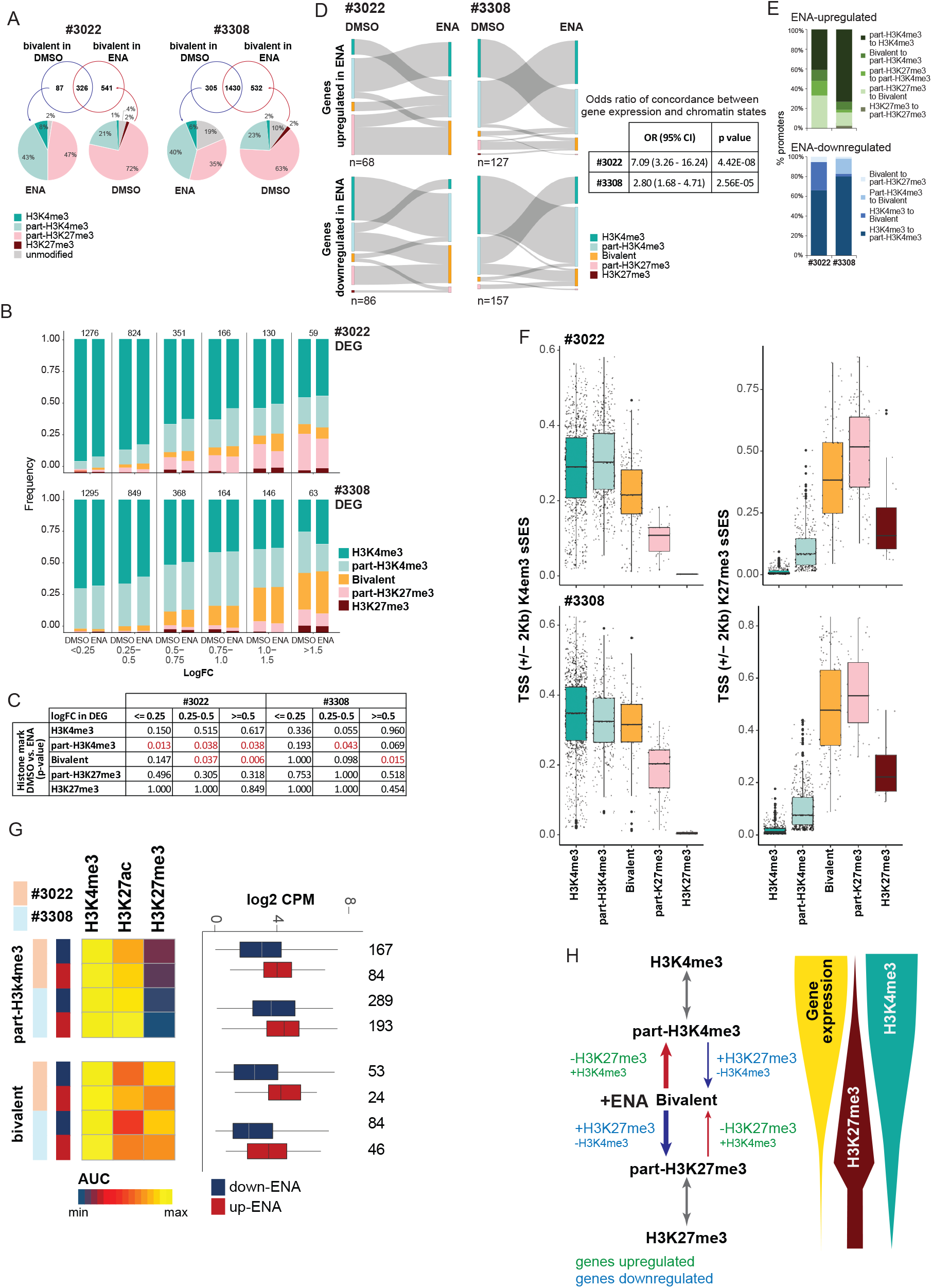
Active and repressive histone marking of DEG of Enasidenib-treated AML progenitors. A) Venn diagrams show bivalent promoters in paired DMSO and ENA-treated samples from day 5 data in #3022 and #3308. Those present under only DMSO or only ENA conditions were analysed for chromatin states in the corresponding ENA and DMSO-treated sample respectively. Histone states are shown in pie charts. The blue arrows indicate the histone states *to* which genes which are bivalent in DMSO only transition, the red arrows indicate the histone states *from* which genes which are bivalent in ENA only transition. B) Bar graphs showing DEGs that were divided according to fold-change (logFC) between DMSO and ENA conditions in #3022 and #3308 (numbers of DEGs are shown above the bars), annotated with chromatin states. C) Analysis of DEGs from (B), grouped as less (logFC </=0.25) or more dynamic (logFC>0.25), comparing frequency of different chromatin states between DMSO and ENA. ENA had a statistically significant (Fisher ’s exact test) higher proportion of chromatin marks in those p-values highlighted in red. D) Sankey plots showing promoters of DEG (numbers of genes shown) which had change of chromatin states between DMSO and ENA conditions in #3022 and #3308. The odds ratio and p-values for concordance between the direction of chromatin change with that of gene expression is shown. E) Bar graphs showing the chromatin state of promoters of concordant ENA-upregulated and downregulated genes in #3022 and #3308. F) Boxplots showing H3K4me3 and H3K27me3 sSES values of gene promoters annotated as monovalent, bivalent or part-bivalent from ENA-treated #3022 and #3308 samples. (Boxplots represent 25-75^th^ quartiles, with a median line; whiskers are 1.5x interquartile range). G) Boxplots show expression (Log2CPM) of ENA-induced up or downregulated genes (from bulk RNAseq data) whose promoters are marked as part-H3K4me3 or bivalent (see Supplementary Figure 3D). The number of genes is included. Alongside, the heatmap is of area under the curve (AUC) values derived from density plots of sSES, showing relative levels of H3K4me3, H3K27ac and H3K27me3 of these genes. H) Our model proposes that ENA reverses the effects of d-2HG in mIDH2 L-Prog by modulating gene expression through shifts to and from the bivalent state. Arrows indicate changes in permissive and repressive histone marks, with the size of the arrow and font indicating dominant pattern of change. Red and blue denote genes up or downregulated by ENA respectively.

### Enasidenib promotes bivalent chromatin states in dynamically expressed genes during differentiation of AML progenitors

Given the effects of d-2HG on TET2 and Jumonji-KDMs (J-KDMs), we studied CpG and histone methylation in ENA-induced differentiation. Using reduced-representation methylation sequencing which enriches for CpG islands (CpGi) in 2 responsive samples (#3022/#3308^RES^) at days (D)2 and D5 of IVC, we found that 4-9% and 1-2% were hypomethylated in ENA and DMSO respectively (Supplementary Figure 4A). To assess methylation and differentially gene expression, we compared 666 DEGs between DMSO/ENA samples from day (D)1-5 and 4044/4548 CpGi (in #3022/#3308^RES^ respectively) represented at both D2 and D5 (Supplementary Figure 4B). Despite ENA-induced hypomethylation at D2, there were no significant DEGs up to this timepoint (Supplementary Figure 4C). At D5, promoters of ENA-upregulated genes were more likely to be hypomethylated than non-DEG (OR 1.2-1.4, p=0.000048-0.008). However, CpGi hypomethylation in upregulated genes was small (mean delta beta: -0.039/-0.075, Supplementary Figure 4B) and there was no association between hypermethylation and downregulated genes. While we cannot rule out a role for CpG hypomethylation in contributing to some gene upregulation, our evidence suggests that other mechanisms are important.

Despite reports of demethylation of multiple histone-3 lysine (H3K) marks by mIDHi^10^, little is known about chromatin changes during differentiation of primary AML progenitors. Bivalent chromatin with both H3K4me3/H3K27me3 marks is poised to be activated or repressed, and modulates gene expression during cell differentiation^17^. To study if ENA restores differentiation by altering bivalent chromatin in a gene-specific manner, we performed chromatin immunoprecipitation sequencing (ChIP-seq)^18^ of H3K4me3, H3K27me3 and H3K27ac in DMSO/ ENA-treated L-Prog harvested at day (D)2 and/ or 5, in 3 responders (#3018/#3022/#3308^RES^) and 2 non-responders (#2917/#2814^NR^).

Consistent with canonical models, H3K27ac and H3K4me3 marks correlated with highly expressed genes from our RNAseq dataset and H3K27me3 correlated with repressed genes (Supplementary Figure 4D). Using a bivalency prediction model^19^, we annotated genomic bins as monovalent (i.e. either H3K4me3 or H3K27me3 only), bivalent, partial (part-) bivalent or unmodified, focusing our analysis on gene promoters. Relationships between monovalent and bivalent marks and expression^17^ were preserved with part-H3K4me3 or part-H3K27me3 genes having intermediate high or low expression respectively. However, ENA treatment did not alter gene expression levels associated with each histone state (Supplementary Figure 4E). Instead, ENA increased promoter bivalent and part-H3K4me3 marks in responders, but not in non-responders (Supplementary Figure 4G). Monovalent H3K4me3 was unchanged by ENA. Furthermore, ENA treatment led to promoters adopting a more permissive histone state, for example, 46-51% gene promoters which were bivalent with DMSO only became part-H3K4me3 or monovalent H3K4me3, and 65-74% of part-H3K27me3 or monovalent H3K27me3 promoters with DMSO became bivalent (Figure 4A).

Next, we studied how ENA-induced chromatin changes relate to gene expression from bulk RNAseq data in responsive samples. ENA-treated cells have a significantly higher proportion of bivalent and part-H3K4me3 marked promoters than DMSO, particularly in the most dynamic DEGs (Figure 4B-C). 13.3% of up, and 21.4% of downregulated dynamic (logFC>0.25) DEG promoters had altered chromatin states between DMSO and ENA (Figure 4D, Supplementary Table S10) and there was significant concordance between direction of chromatin change and gene expression (OR=2.80-7.09 Figure 4D-E). To model the most likely (de)methylation events here, we computed scaled sum of enrichment scores (sSES) for promoter marks (Figure 4F). 67%-84% of concordant ENA-upregulated genes transition from bivalent to more permissive histone states. These are at least 2.9x more likely to lose H3K27me3 than gain H3K4me3 (Supplementary Table S11a). >95% of downregulated gene promoters move from activated states towards bivalency, mainly through gain of H3K27me3 marks (Figure 4E-F). These are at least 4x (4.28-11.03) more enriched with polycomb repressor complex 2 (PRC2)/EZH2 targets than randomly selected genes, and are not enriched for PRC1/BMI1 targets (Supplementary Tables S11b-c).

In bivalent marked genes, those upregulated by ENA (including *CEBPA, AZU1, ELANE* and *PRTN3*) have fewer H3K27me3 and more H3K27ac marks, compared with downregulated genes (such as *GATA2*, Figure 4G, Supplementary Figure 4I, Supplementary Tables S12a-b) while H3K4me3 levels remain unchanged. Our model is therefore one where ENA-induced dynamic gene expression is regulated by transitions between bivalent and part-bivalent states with a continuum of methylation and demethylation of H3K27 during gene activation rather than of H3K4 (Figure 4H). This also reflects preferential inhibition by d-2HG of KDM4A/B and KDM6A/B which demethylate bi- and/or tri-methylated H3K9/36/27 over the H3K4me3 demethylase KDM5B^8^.

### Differentiation of AML progenitors is driven been increased chromatin accessibility of TF targets

Transitions in H3K4me3/ H3K27me3 states in our data from less, to greater transcriptional permissiveness is correlated with increases in promoter H3K27ac (Supplementary Figure 4D and 5A). Accessible chromatin in ATAC-seq positively correlates with H3K27ac, transcriptionally active gene promoters, enhancers and transcribed regions^20^, and anti-correlates with H3K27me3^21^. To improve resolution of ENA-induced chromatin changes in the continuum of differentiation, and to employ an orthogonal method to study chromatin, we performed single-cell (sc)ATAC-seq in two responder (#3018/#3022^RES^) and two non-responder (#2917/#2755^NR^) cells harvested at D5 to capture diverse cell types.

Using ArchR^22^, we clustered cells based on TF motif accessibility and integrated scRNA-seq and scATAC-seq data to derive a gene integration matrix. As with our scRNAseq analysis, we excluded CD14-expressing monocytes to focus our analysis on neutrophil differentiation (Supplementary Figure 5B). We constructed pseudotime trajectories using *ELANE* expression, which for descriptive purposes were divided into early, mid and late pseudotime phases (Figure 5A, Supplementary Figure 5C). Compared with DMSO, ENA increased accessibility of peaks across all phases (Supplementary Figure 5D). Peaks which were more accessible in ENA were closer to transcription start sites (TSS) compared with DMSO (Figure 5B), suggesting that ENA targets promoter accessibility. TF motif usage during ENA-induced differentiation is concordant with engagement of regulons from scRNAseq. ENA-treated cells progress beyond engagement of AP-1 (JUN/FOS) motifs, to CEBP motifs, while DMSO-treated cells have persistently high enrichment of GATA, core binding factor (CBF/RUNX) and AP-1 motifs but fail to engage CEBP (Figure 5C). In ENA-treated cells, footprinting analysis of scATACseq showed deepest GATA2 footprints in undifferentiated (U) cell clusters, early and late GM cells. The reverse is true for CEBPA and CEBPE footprints which are deepest in late GM cell clusters (Figure 5D).

**Figure 5.**
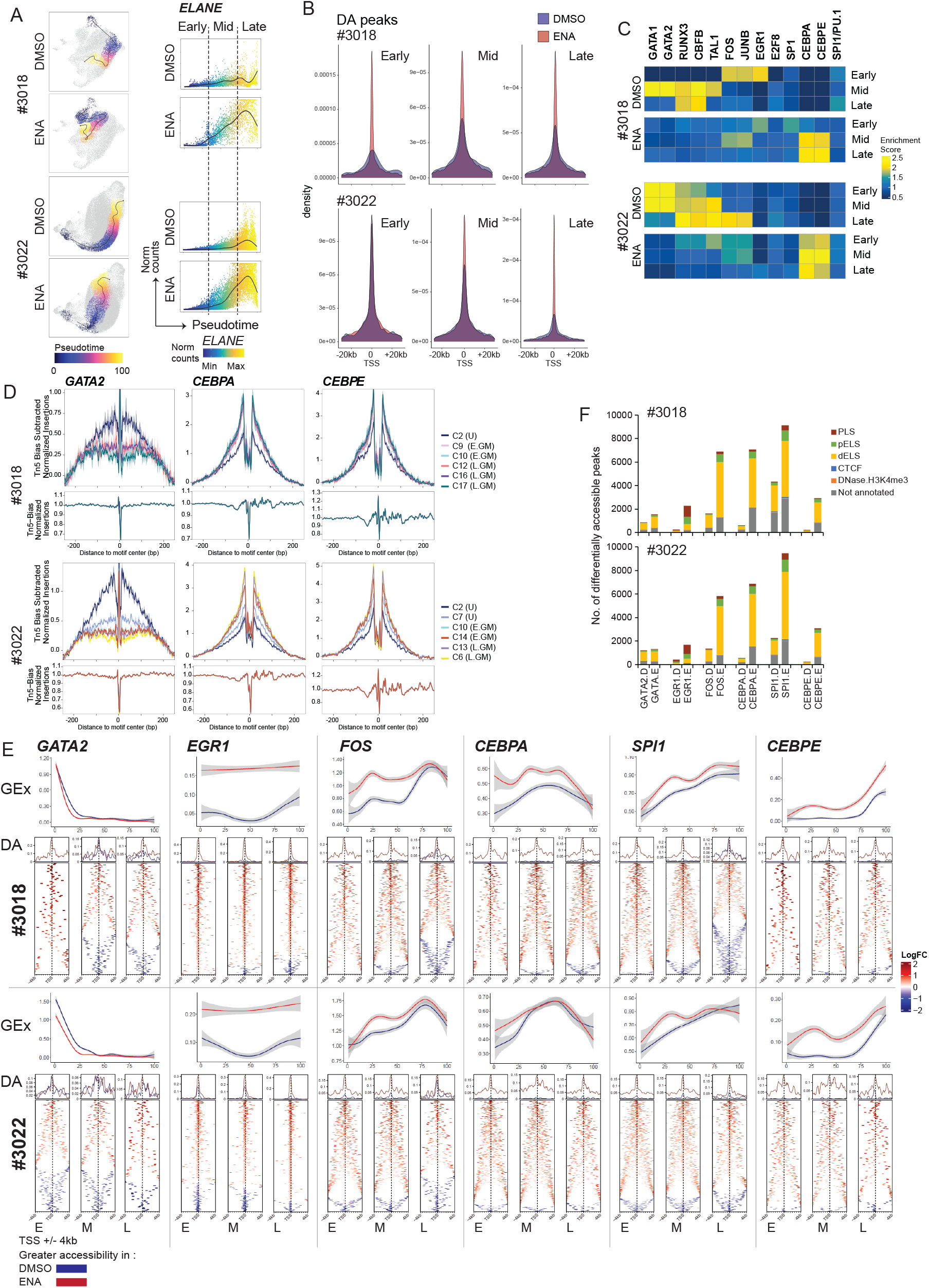
ENA induces chromatin accessibility at promoter regions of transcription factor gene targets. A) Paths of pseudotime trajectory of DMSO and ENA projected onto UMAP clustering of scATACseq cell clusters in responsive samples #3018 and #3022: The ’start ‘ (see Supplementary Figure 7B) and ‘end ‘ clusters were chosen based on *ELANE* expression, which is also shown as a line graph (right panel). Pseudotime for these cells were divided into three equal early, mid and late bins for ease of description. B) Graphs show mapping of differentially accessible peaks to genomic regions relative to transcription start sites (TSS) +/- 20kb in early, mid and late pseudotime, in DMSO or ENA treated cells from #3018 and #3022. C) TFs motif (labelled by CIS-BP annotation) enrichment analysis of DA peaks in early, mid and late phases of pseudotiome trajectory in DMSO and ENA treated cells from #3018 and #3022, of GATA, core binding factor, EGR1, AP-1 (JUN/FOS) and CEBP family TF motifs. The scale shows relative enrichment per motif, and per patient. D) Footprinting analysis for GATA2, CEBPA and CEBPE motifs in ENA-treated clusters enriched for undifferentiated (U), early GM (E.GM) and late GM (L.GM) cells, from #3018 and #3022. E) Gene expression from scRNAseq data of #3018 and #3022 (upper panels) is shown alongside stacked heatmaps of DA peaks harbouring motifs for *GATA2, EGR1, JUNB, FOS, CEBPA, SPI1/PU*.*1/PUl1* and *CEBPE* within 4kb of the TSS of putative target genes in early (E), mid (M) and late (L) trajectory phases of #3018 and #3022. Positive and negative log-fold change (logFC) values are peaks more accessible in ENA and DMSO respectively. The dotted line denotes the TSS. The summary plot is also shown (blue line= DMSO, red line= ENA). F) Bar graphs of numbers of differentialluy accesible peaks with TF motifs in DMSO and ENA-treated cells from #3018 and #3022. Peaks are also annotated with Encode candidate cis regulatory elements (cCREs, PLS, proximal promoter-like signature, pELS and dELS proximal and distal enhancer-like signature respectively, CTCF, CTCF-only and DNAase-H3K4me3, promoter-like signature not within 200bp of a TSS).

We validated TF binding sites (TFBS) from scATACseq data using published ChIP-seq datasets (Supplementary Table S13). Most TFBS were markedly increased and in ENA-treated cells (Figure 5E). This suggests that ENA induces target accessibility in addition to upregulation of pro-differentiation TFs like EGR1, AP-1, CEBPA/E and SPI1 to drive differentiation. Furthermore, using Encode^23^ annotation, ENA-stimulated peaks were more frequently proximal cis regulator elements (CREs) (i.e. promoter-like signature, PLS, and proximal enhancer-like signature, pELS; ENA vs. DMSO, #3018: 27.2% vs. 19.0%, #3022: 28.15% vs. 25.2%, Wilcoxon signed-rank test, p=0.0068, Figure 5F). However, compared with pro-differentiation TFs, ENA ’s effects on GATA2 targets were less marked, with less enrichment for proximal CREs (Figure 5F).

### Regulation of expression of key transcription factors in differentiation through differential binding and accessibility of promoter and *cis* regulatory elements

Co-engagement of multiple TF regulons or networks suggests cross-regulation between stem-progenitor and pro-differentiation TFs. Thus, we constructed regulatory networks using CRE accessibility, motif usage and gene expression from responsive samples (#3018/#3022^RES^). Almost all interactions originating from pro-differentiation TFs are greater with ENA (pink-red arrows, Figure 6A), while those which are more prominent with DMSO tend to originate from stem-progenitor TFs (blue arrows, Figure 6A).

**Figure 6.**
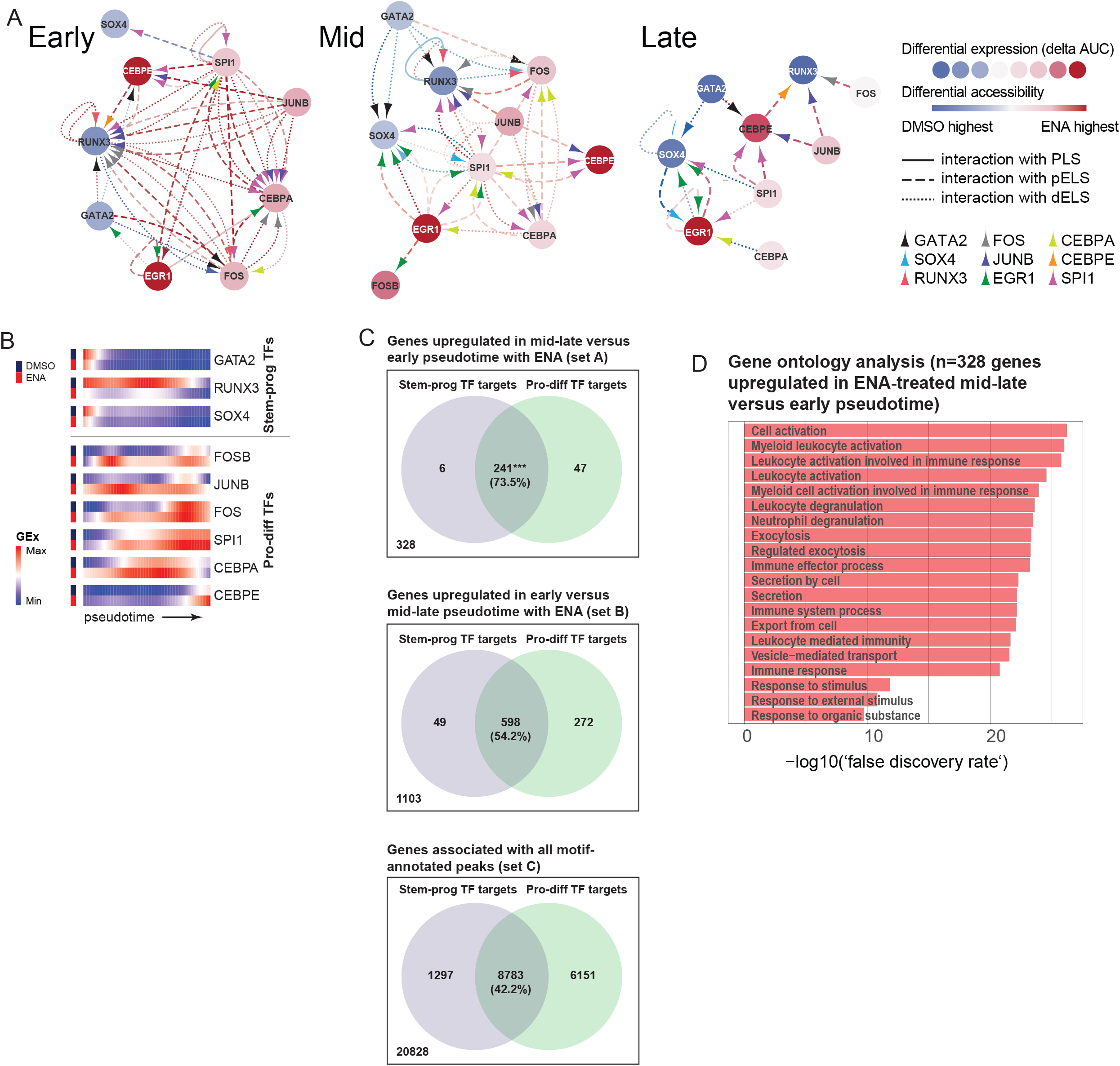
Active transcription factor regulatory networks during differentiation of AML progenitors. A) Regulatory networks showing relationships between TFs during early, mid and late phases of differentiation were constructed using data from scATACseq, with gene expression data from scRNAseq used to infer activating or repressive effects. Nodes are coloured according to relative gene expression level (delta AUC, blue: highest in DMSO, red: highest in ENA). Arrowheads are colour-coded by TF shown in the key. Colours of connecting lines indicate increased accessibility (blue: highest in DMSO, red: highest in ENA), whereas solid or dashed lines indicate putative CRE interactions between TFs. B) Heatmaps of expression (GEx) of stem-progenitor (stem-prog) and pro-differentiation (pro-diff) TFs in DMSO and ENA treated conditions. The data is scaled by minimum-maximum in each gene and are the average of scRNAseq data from all 4 responsive patients (as in Figure 4E, which showed regulon expression). C) Venn diagrams showing overlap of target genes of stem-progenitor (stem-prog) and pro-differentiation (pro-diff) TFs from : (upper panel, set A) a set of genes upregulated in ENA-treated mid-late versus early pseudotime, (middle panel, set B) genes upregulated in ENA-treated early versus mid-late pseudotime and (lower panel, set C) genes selected only on the basis of annotation with a known TF motif from our scATACseq data. Differential gene expression was obtained from combined scRNAseq data from 4 responsive patients. *** indicates significantly increased proportion of dual-regulated genes with p values of <0.0001 from Fisher exact (set A versus set B) or chi-squared test (set A versus set C). D) Top 20 terms from gene ontology analysis of 328 genes (set A, upregulated in ENA-treated mid-late versus early pseudotime).

In early ENA-induced differentiation, stem-progenitor TFs are targeted by pro-differentiation TFs, e.g. putative EGR1 binding to a distal enhancer-like signature (dELS) of GATA2, and engagement of RUNX3 CREs by AP-1, CEBPA/E and SPI1 (Figure 6A, Supplementary Figure 6A-B). Expression patterns suggests that stem-progenitor TFs are repressed by pro-differentiation TFs, but there are also auto-regulatory interactions, e.g. in RUNX3 (Figure 6A-B). Targeting of CEBPA and CEBPE by EGR1, FOS, JUNB and SPI1 is also enhanced by ENA from early phase onwards and represent events which initiate AML progenitor differentiation (Figure 6A, Supplementary Figure 6C-D). With CEBPA, we observe increased accessibility of +4/21/29/34 enhancers^24^, and predict positive regulation by EGR1, JUNB, FOS and SPI1 (Supplementary Figure 6C). This is consistent with reports in mouse models^25^ and a megakaryocytic-erythroid G1ME cell line^26^ that *Cebpa* is regulated by Egr1 and Spi1. We also detect a putative repressive interaction between GATA2 and a *CEBPE* dELS which is increased in DMSO in early phase (Supplementary Figure 6D). Conversely, in mid and late phases, expression of stem-progenitor TFs is decreasing (Figure 6B) and there is increased repressive targeting of SOX4 and RUNX3 by EGR1, AP-1 and SPI1 in ENA-treated cells (Figure 6A, Supplementary Figure 6B).

### Neutrophilic genes are dual regulated by stem-progenitor and pro-differentiation transcription factors in AML progenitors

Our bulk and scRNAseq data indicate that ENA has little effect on transcription when stem-progenitor TFs are most highly expressed in early differentiation (Figure 6B), even though we observe increased in accessibility of stem-progenitor TF targets at this time. We reasoned that this could reflect the repressive effect of stem-progenitor TFs on neutrophilic differentiation genes and examined our data for targets of both stem-progenitor and pro-differentiation TFs. We identified 328 genes which with ENA, were lowly expressed in early pseudotime, and then upregulated in mid-late pseudotime (Figure 6C). These are enriched for myeloid and neutrophil genes (Figure 6D), and 241 (73.5%) genes were targets of both stem-progenitor and pro-differentiation TFs, indicating dual regulation (Fisher ’s exact test p<0.0001, compared with unselected genes or those upregulated in early pseudotime, Figure 6C). These included pro-differentiation TFs *CEBPE, FOS* and *SPI1*, and neutrophil effector genes *CSF3R, FCGR1A, MNDA, MPO* and *SRGN*. Together, these data indicates that stem-progenitor TFs repress neutrophil differentiation genes, and that ENA induces pro-differentiation TFs to switch off stem-progenitor programmes to restore differentiation.

## DISCUSSION

Differentiation arrest in IDH-mutant tumours is vulnerable to therapeutic targeting. In AML, compared with normal GMP, arrested IDH2-mutant progenitors downregulate GM genes while upregulating stem-progenitor genes, and have reduced expression of cell cycle (CC) genes. We hypothesised that restoration of normal GM cell fate involves at least the downregulation of stem-progenitor pathways, and engagement of CC and GM programmes. D-2HG produced by mutant IDH suppresses DNA CpG methylation via TET2, and histone lysine demethylation via Jumonji-KDMs. However, the link between epigenetic changes and gene expression has not been demonstrated in primary human IDH-mutant AML progenitors.

Using an *in vitro* differentiation assay which mirrors clinical observations *in vivo*^4^, we find in single cell transcriptomic trajectories that ENA induces sequential downregulation of stem-progenitor programmes followed by simultaneous expression of CC and GM programmes in the same cells to restore differentiation. We observed rapid downregulated GATA2/RUNX3/SOX4^R^ (GRS^R^) during differentiation, with sustained RUNX3^R^ expression being associated with non-response to ENA. Crucially, there is sustained co-expression of early pro-differentiation regulons (EGR1/JUN/FOS^R^) in the same GRS^R^-positive cells, demonstrating for the first time, an early transitional subpopulation that marks a turning point where AML progenitors embark on a differentiation fate.

*GATA2, RUNX3* and *SOX4* are implicated in leukaemogenesis and, to our knowledge, this is first evidence of their co-expression in arrested primary AML progenitors, and their downregulation during drug-induced differentiation. Haploinsufficiency of *GATA2* causes of familial AML and MonoMac syndrome and its enhancer ectopically activates *EVI1* in inv(3)/t(3;3) AML^27^. *GATA2* is upregulated in IDH2-mutant AML progenitors relative to normal GMP and is more rapidly downregulated on ENA-induced differentiation than controls. This is different from observations in mice where, in an *Idh2/Flt3*^*ITD*^ AML model, *Gata2* upregulation is associated with response to combined IDH2 and FLT3 inhibition^28^. Paradoxically, *Gata2* is also downregulated in mouse granulocyte-monocyte-committed progenitors compared immature myeloid progenitors^29^. We were hampered by the lack of suitable IDH2-mutant AML cell line models that undergo neutrophil differentiation with IDH2 inhibitors which prevented mechanistic demonstration of the regulatory relationship between *EGR1* and *GATA2*. However, we do observe ENA-induced increased accessibility of a known *EGR1* binding site at a CRE - 5kb from the *GATA2* promoter. *Sox4* is a target of *Cebpa* in mouse HSPCs and its overexpression due to *Cebpa* inactivation is leukaemogenic^30^. While we did not observe direct interaction between *CEBPA* and *SOX4*, we do have evidence of negative regulation of *SOX4* by *EGR1*. Furthermore, *SOX4* regulon (SOX4^R^) expression is increased in non-responsive versus responsive AML progenitors. A similar but more marked difference is also seen with RUNX3^R^. *RUNX3* expression is increased in some AMLs where it is a marker of poor prognosis^31^. Its overexpression in normal HSPCs has recently been reported to inhibit GM and erythroid differentiation and drive expression of lymphoid programmes in AML cells ^32, 33^. Mirroring leukocytosis commonly observed in ENA-treated patients, *in vitro* we found that ENA stimulated proliferation of late GM cells underpinned by co-expression of cycling and GM pro-differentiation programmes. The subversion of normal differentiation programmes by IDH mutations has also been noted in an *Idh1*-mutant mouse model where hepatocyte differentiation arrest is associated with reduced expression of targets of Hepatocyte Nuclear Factor-4α, a regulator of hepatocyte quiescence and lineage specification ^2^.

From scATACseq motif analysis, we confirmed downregulation of stem-progenitor and engagement of pro-differentiation programmes during differentiation. In our model of regulatory interactions, *GATA2, SOX4* and *RUNX3* were downregulated by pro-differentiation TFs, consistent with co-expression of the same regulons in early transitional cells from scRNAseq data. The same model identified cross-regulation of *EGR1*/AP-1 TFs and *SPI1, CEBPA* and *CEBPE* which is also consistent with regulon co-expression during mid-late phases of differentiation. The role of *CEBPA* during ENA-induced differentiation is also underlined by the finding that *CEBPA* mutations^34^ were associated with failure to respond to mIDHi therapy and is a mechanism for disease relapse^35, 36^.

We show a close temporal relationship between histone methylation and chromatin accessibility with differential gene expression with ENA-induced differentiation. We demonstrate for the first time in primary AML progenitors that ENA-induced gene expression is associated with modulation of bivalent and part-bivalent chromatin states, through H3K27me3 demethylation (mediated by KDM4/6) of GM-differentiation genes. Of note, a recent report of mouse granulo-monocytic differentiation using an unbiased CRISPR-Cas9 perturbation screen found that *Kdm6a* promotes myeloid versus megakaryocytic-erythroid fate linage priming ^37^. Our analysis indicates that downregulated genes including *GATA2* gain H3K27me3 via PRC2/EZH2 activity. While PRC2 is implicated in leukaemogenesis e.g. in mixed lineage leukaemia models^38^, in the context of IDH-mutant AML, it may act as a tumour suppressor. Interestingly, consistent with our observations, a recent report showed that EZH2 knockdown in IDH1-mutant TF-1 AML cells promotes cytokine-independent growth ^39^ although the mechanism by which this is regulated by d-2HG is unknown. However, while we found that loss of H3K4me3 (mediated by KDM5) occurs less frequently than gain of H3K27me3 in ENA-downregulated genes, this report implicated KDM5 in IDH1-mutant transformation in glioma and AML. This may indicate potential differences between IDH1 and IDH2-mutant disease, and varying roles of IDH mutations in different tumour types ^39^.

Loss of repressive histone marks in bivalent and part-bivalent chromatin correlates with the increased accessibility in response to ENA. Using scATACseq we found increased usage of pro-differentiation TF motifs, related to increased expression and target accessibility of those TFs. Furthermore, ENA induced increased accessibility of targets of *SPI1* and *CEBPE* in early differentiation, when expression levels of these TFs were low. This may be a priming mechanism, for example, accessibility of *GATA1* targets were found to precede subsequent *GATA1* expression in megakaryotic-erythroid progenitors^40^. Importantly, we found that the majority of genes upregulated by ENA during established differentiation are dual-regulated: repressed by stem-progenitor TFs initially, and then activated by pro-differentiation TFs.

Our model of how d-2HG transforms normal into arrested leukaemic progenitors and the events that lift differentiation blockade, highlight co-expression of conflicting progenitor-sustaining and pro-differentiation transcriptional programmes occurring at a critical junction in haematopoiesis that ultimately permits terminal differentiation. Our observations also place modulation of chromatin accessibility via KDM4/6 and PRC2 at centre-stage. For the first time, we show that these chromatin changes are associated with dynamic transcriptional programmes that initiate and sustain neutrophil differentiation from primary AML progenitors, and identify programmes associated with treatment resistance. Our findings in primary patient samples help to clarify previous conflicting observations on the IDH-mutation driven mechanisms in haematopoiesis. Crucially, the finding that IDH2 inhibitors appear to work through Jumonji-KDM pathways provides a rationale for synergism underlying increased efficacy in combining IDH inhibitors with DNA hypomethylating agents^41^.

## Data and code sharing statement

RNAseq Data from normal bone marrow populations were accessed from ArrayExpress: E-MTAB-2672 and E-MTAB-5456. High-throughput data (RNAseq, ChIPseq, Single-cell RNAseq, and single-cell ATACseq) are available for download from the King ’s Open Research Data System (KORDS) - DOI: 10.18742/20766250. Analysis codes can be made available upon request to the corresponding author.

## Acknowledgments

we thank Benjamin Brown, Anna Corby and Mattia Marinucci for their technical assistance in this work. This study and L.Q. and P.C. were funded by UKRI/ MRC grant MR/R007608/1.

## Authorship

L.Q. conceived the study, designed and performed the experiments, interpreted the data, wrote and reviewed the manuscript; D.R.A.S. performed bioinformatic and statistical analyses, performed experiments, interpreted the data, wrote and reviewed the manuscript; P.C. and P.C.A. performed experiments, interpreted the data, wrote and reviewed the manuscript; S.M., R.H., O.Y., P.S.Z, and S.K., performed experiments, interpreted the data, and reviewed the manuscript; A.S., M.M., B.U, V.P.L, C.X.S., A.K.G., M.H., P.V., and T.A.M provided data and reviewed the manuscript. All authors approved the submission version.

