## Supplemental figures for "Dysregulation of chromatin via H3K27 methylation underpins differentiation arrest in Isocitrate dehydrogenase-mutant Acute Myeloid Leukaemia"

### Supplementary Figure 1

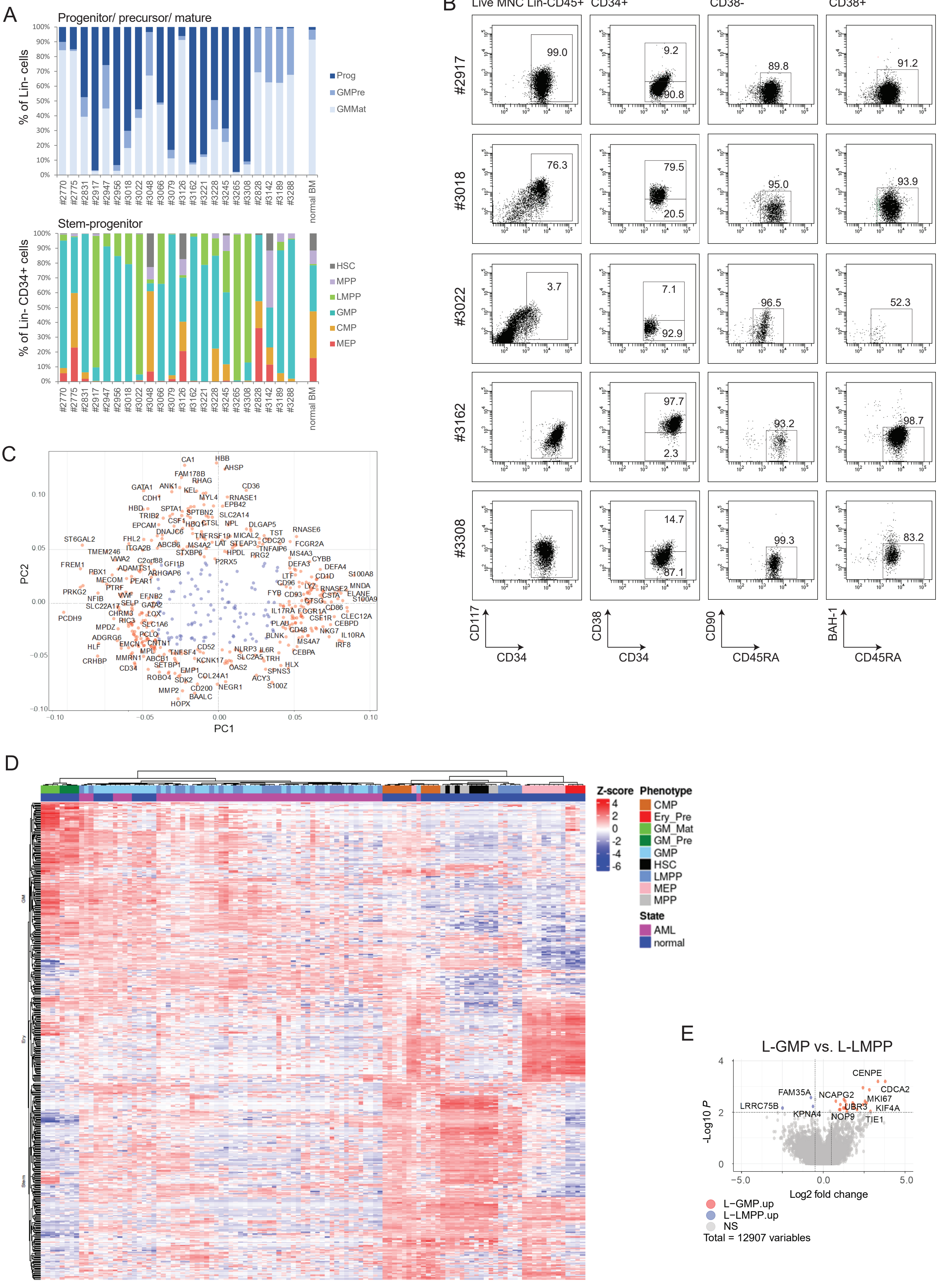

Supplementary Figure 2

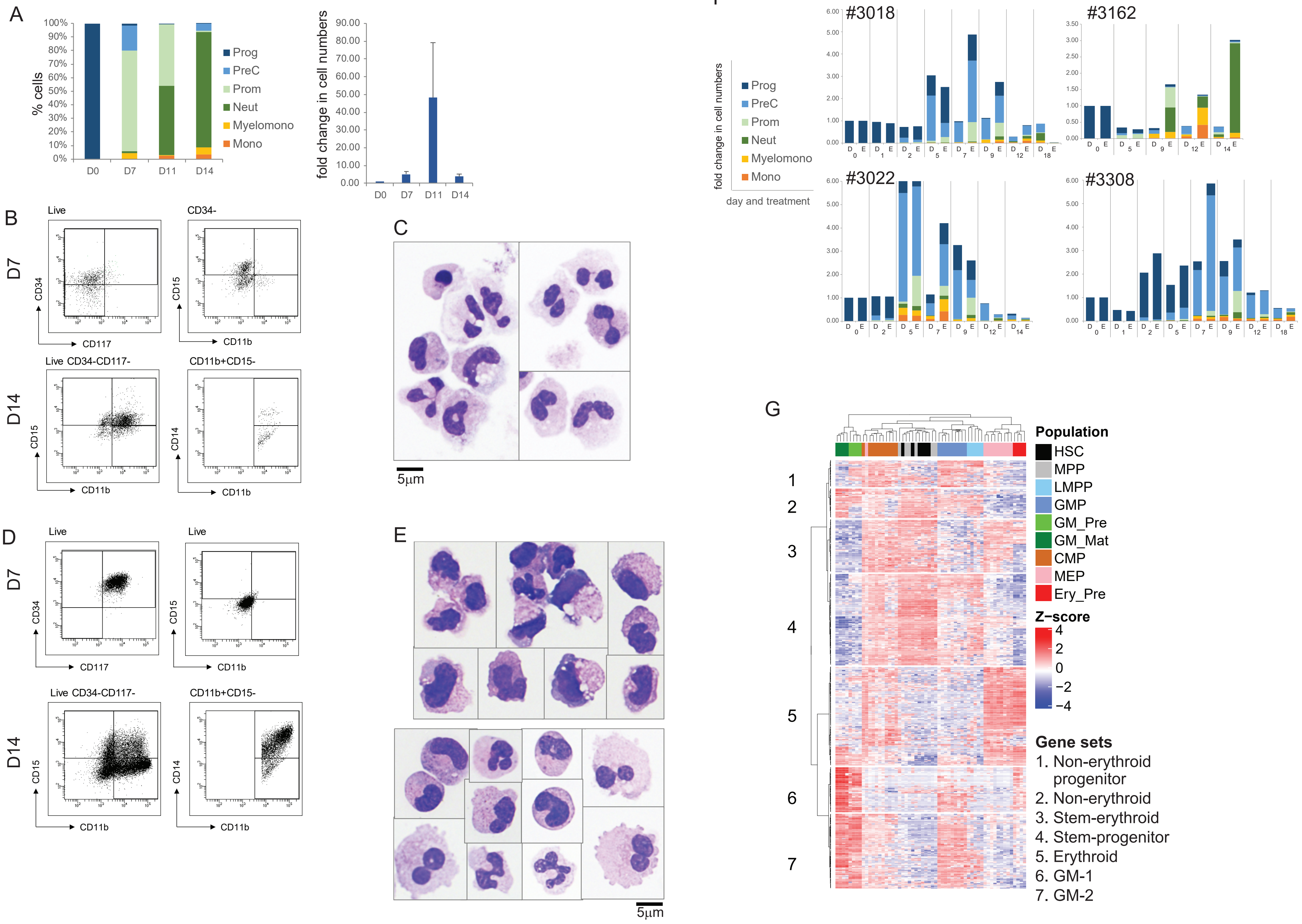

Supplementary Figure 3

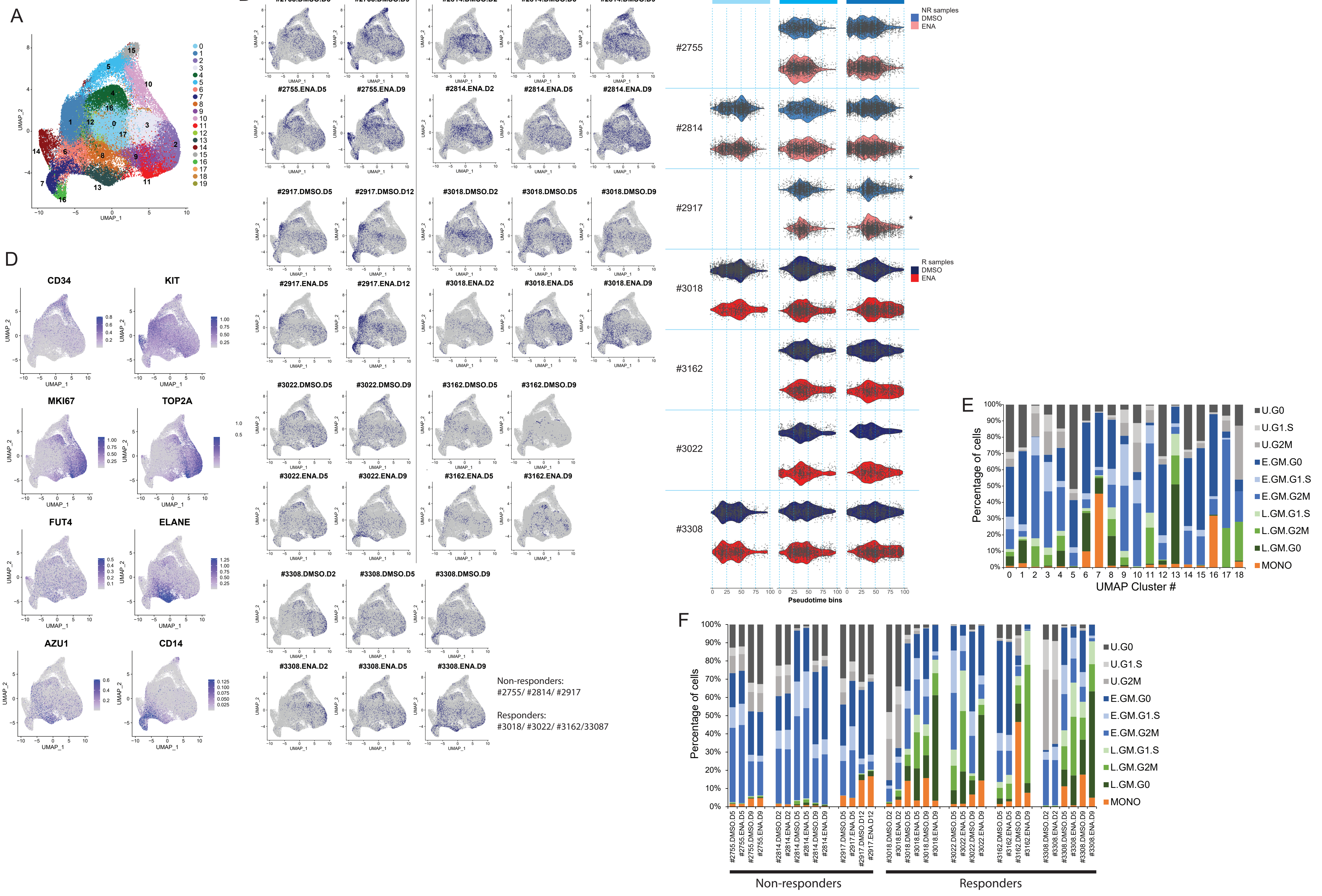

### Supplementary Figure 4

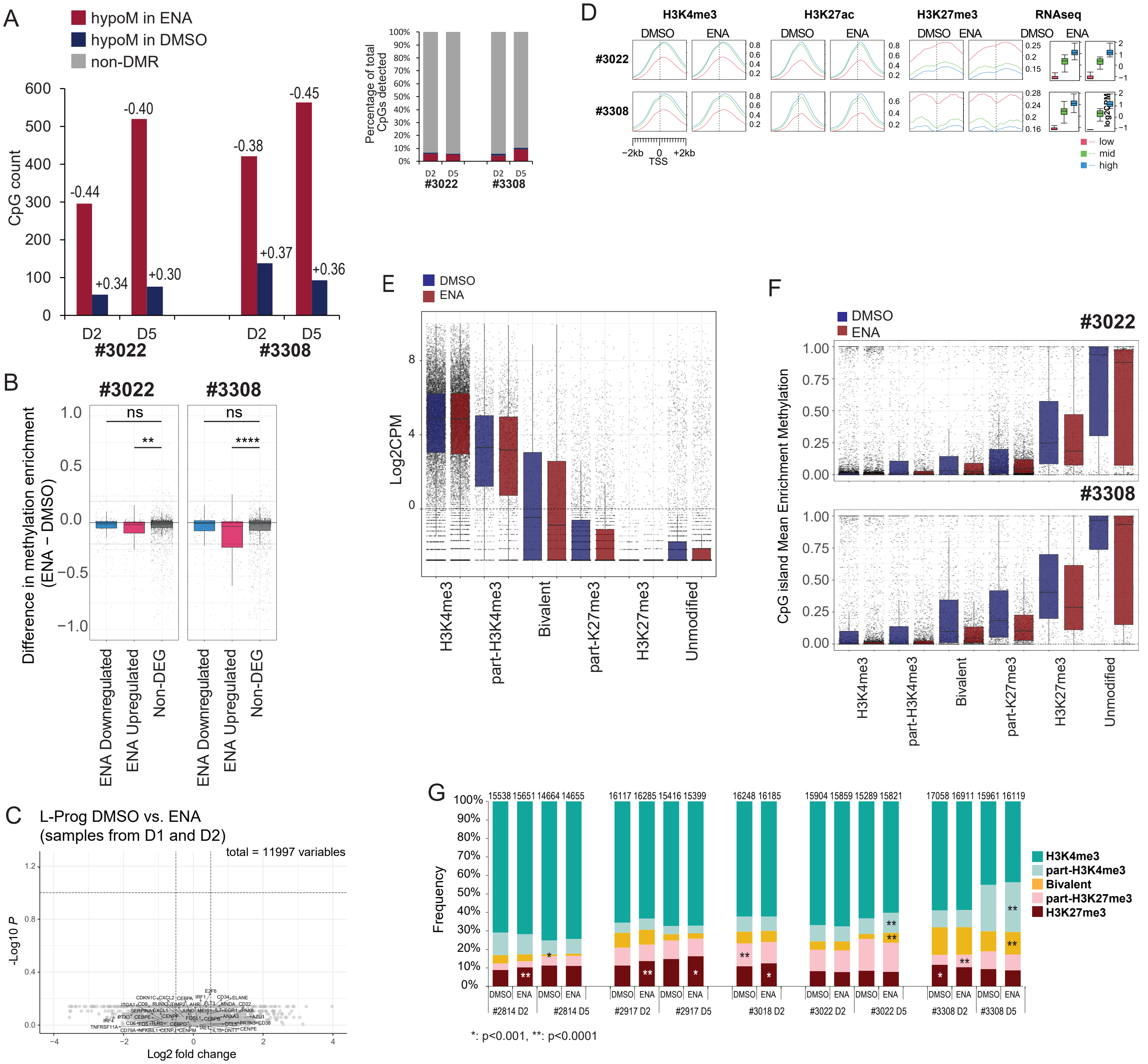


## H

| Genes | #3022 Histone states (ChIPseq data at D5, ENA versus DMSO) | | | #3308 Histone states (ChIPseq data at D5, ENA versus DMSO) | | |
|---|---|---|---|---|---|---|
| **CpGi Methylation (at D2 ENA versus DMSO)** | More permissive | Less permissive | Unchanged | More permissive | Less permissive | Unchanged |
| HypoM in ENA | **13 (5.4%)** | **23 (9.5%)** | **205 (85%)** | **25 (6.4%)** | **48 (12.4%)** | **315 (81.2%)** |
| | OR = 1.15 (0.59 - 2.05) p = 0.63 | OR = 1.86 (1.13 - 2.94) p = 0.008 | OR = 0.63 (0.43 - 0.94) p = 0.02 | OR = 0.64 (0.41 - 0.96) p = 0.03 | OR = 0.91 (0.65 - 1.25) p = 0.64 | OR = 1.29 (0.99 - 1.70) p = 0.055 |
| HyperM in ENA | **5 (10.9%)** | **5 (10.9%)** | **36 (78.2%)** | **14 (11.1%)** | **16(12.7%)** | **96 (76.2%)** |
| | OR = 2.46 (0.75 - 6.31) p = 0.07 | OR = 2.16 (0.66 - 5.53) p = 0.1 | OR = 0.40 (0.19 - 0.92) p = 0.03 | OR = 1.16 (0.61 - 2.05) p = 0.54 | OR = 0.94 (0.51 - 1.60) p = 1 | OR = 0.96 (0.63 - 1.50) p = 0.83 |
| non-differentially M | **204 (4.7%)** | **231 (5.4%)** | **3889 (89.9%)** | **834 (9.7%)** | **1148 (13.4%)** | **6598 (76.9%)** |

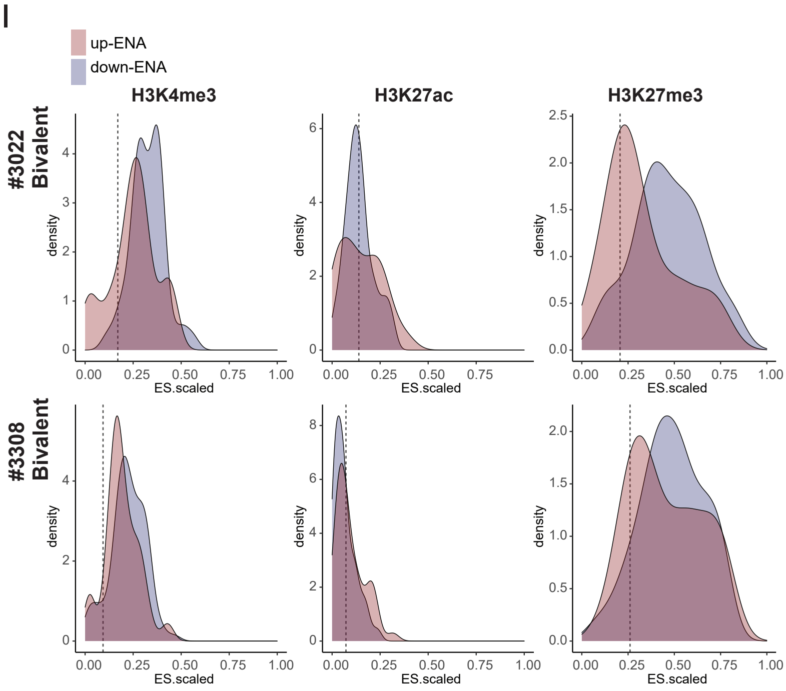


Scaled ES curves shown are for bivalent promoters from #3022 and #3308

Supplementary Figure 5

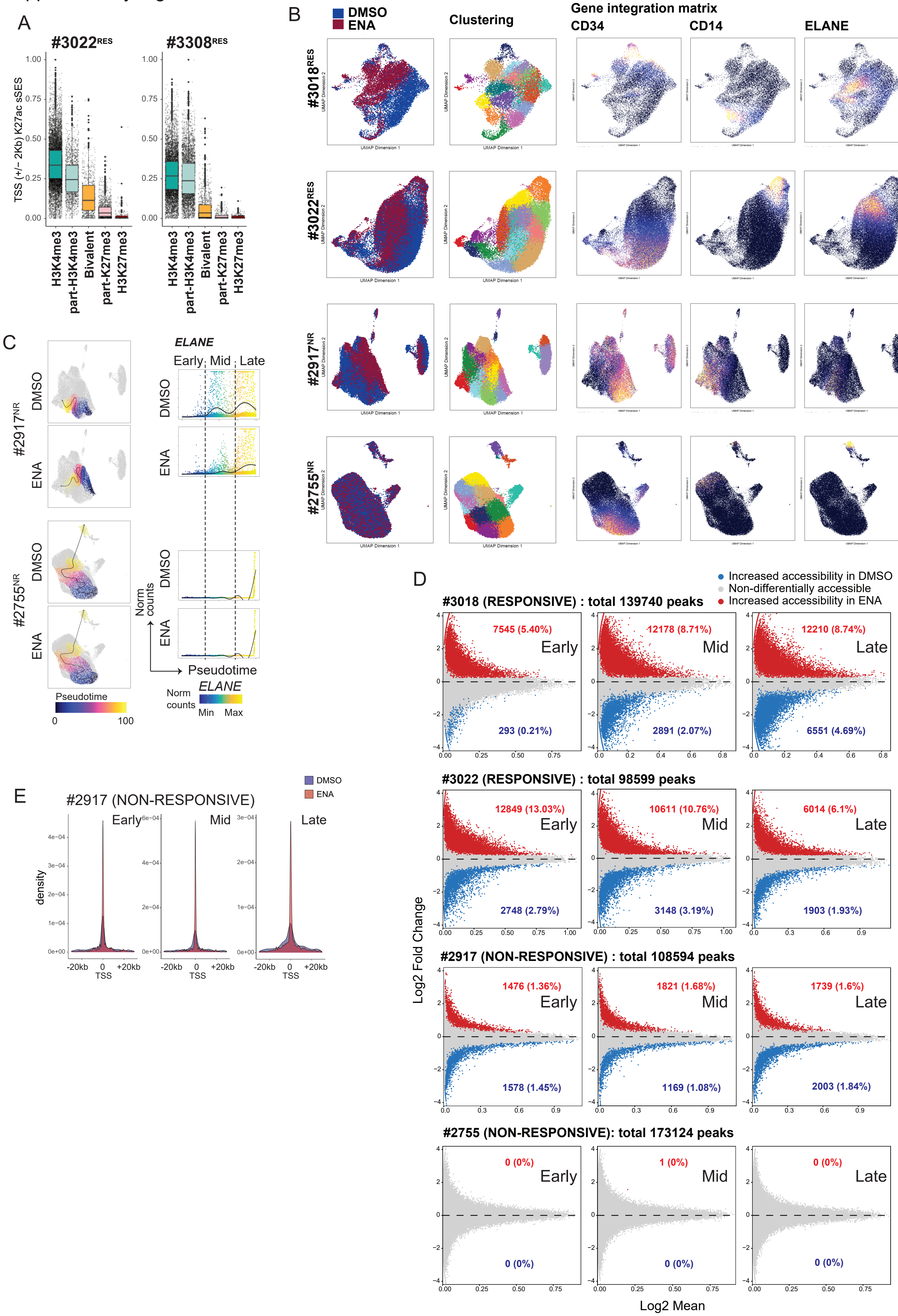

Supplementary Figure 6

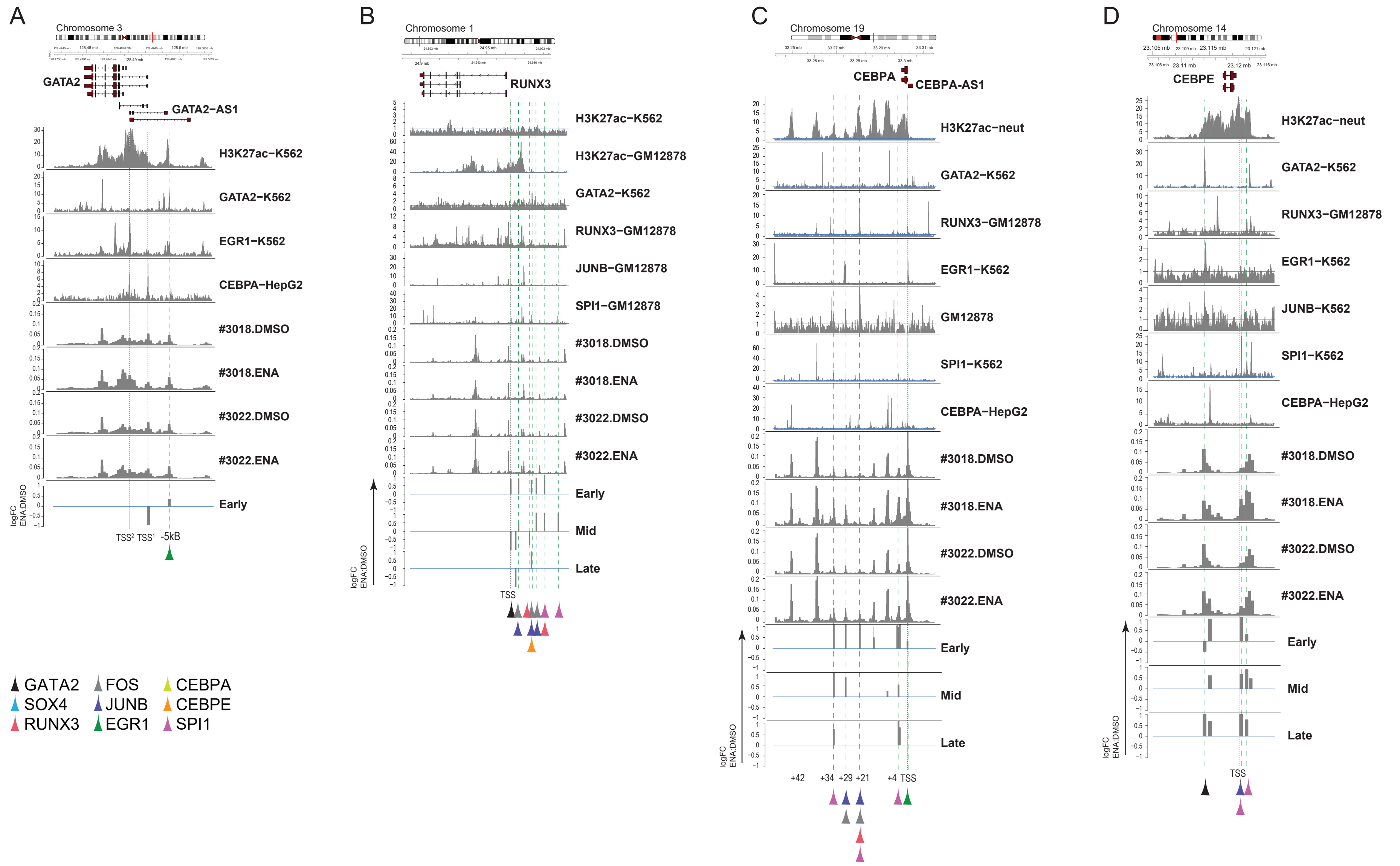
